# PANDA: Protein All-atom Nested-tree Denoising Architecture for End-to-End Generation

**DOI:** 10.64898/2026.09.11.750898

**Authors:** Jie Bai, Huifeng Jiang

## Abstract

Deep generative models have expanded the scope of computational protein design, yet most approaches still separate backbone generation from sequence assignment or rely on latent and torsional representations that do not operate directly on all atoms in Cartesian space. We present PANDA, an end-to-end generative architecture that performs denoising in a unified all-atom representation, recovering sequence identity directly from atomic occupancy patterns. By coupling global and local coordinate tracks and introducing a sampling strategy that preserves side-chain geometry, PANDA achieves the highest self-consistency design success across protein lengths among evaluated all-atom methods. For functional design it conditions on pairwise distances between motif atoms rather than fixing their coordinates, allowing the functional atoms to be positioned jointly with the scaffold. On the Atomic Motif Enzyme benchmark, PANDA preserves motif geometry and delivers substantially higher scaffolding success than previous approaches, particularly for larger motifs. PANDA thus provides an efficient route to high-quality all-atom generative design with flexible atomic-level control.

## Introduction

Deep generative models have substantially expanded the scope of computational protein design. Early approaches commonly separated backbone generation from sequence design. Backbone models such as RFdiffusion (Watson et al., 2023), SCUBA (Chen and Liu, 2024), and Proteina (Geffner et al., 2025b) generate a structural scaffold, after which inverse-folding models such as ProteinMPNN (Dauparas et al., 2022) or ESM-IF1 (Hsu et al., 2022) assign amino acid sequences. This strategy has enabled the generation of stable and structurally diverse proteins and has been extended to systems containing non-protein components. RFdiffusion-AA (Krishna et al., 2024) and LigandMPNN (Dauparas et al., 2025), for example, incorporate nucleic acids, ligands, and cofactors into protein design workflows. However, backbone generation does not explicitly model the side-chain geometry that determines many binding and catalytic interactions. A subsequent sequence-design stage therefore has to recover these atomic arrangements after the backbone has already been generated.

Several methods have moved toward joint structure and sequence generation. Multiflow (Campbell et al., 2024) jointly models amino acid identities and backbone coordinates in a flow-matching framework. Other approaches extend backbone generation to side-chain structure. ProtPardelle (Chu et al., 2024) predicts backbone coordinates together with side-chain rotamer ensembles, while APM (Chen et al., 2025) augments backbone diffusion with torsion-angle prediction. P(all-atom) (Qu et al., 2024) and RFdiffusion3 (Butcher et al., 2025) use AlphaFold3-inspired atom representations in which absent atoms are colocated with C*α*, enabling all-atom generation with a uniform representation. Pallatom-Ligand (Wang et al., 2026) further extends this framework to ligand-conditioned generation. La-Proteina (Geffner et al., 2025a) instead uses a pretrained variational autoencoder to represent side-chain conformations in a continuous latent space, and Proteina-Complexa (Didi et al., 2026) extends this strategy to multi-chain systems and non-protein components. Despite these advances, amino acid identity remains an explicit discrete prediction in these approaches. Consequently, sequence assignment and atomic coordinate generation remain distinct components of the generative process.

We present PANDA, a Protein All-atom Nested-tree Denoising Architecture that represents residue identity through atomic geometry rather than a separate discrete prediction. Prior all-atom models either retain an explicit sequence head or colocate absent atoms with C*α*, which separates identity from the coordinate trajectory or erases residue-specific missing-atom patterns. PANDA instead recovers amino-acid identity from occupancy in a nested atomic tree, implemented with a dual-track denoiser and a sampler matched to the local geometry. Functional motifs are conditioned through pairwise distances rather than frozen coordinates. Together, these components provide an end-to-end framework for all-atom protein generation and functional scaffold design.

## Results

### A nested-tree representation and a global-local coordinate frame unify all-atom denoising

All-atom generation differs from backbone generation because residue identity determines which side-chain atoms should be present. PANDA represents each residue by a nested tree of 28 atomic slots that follow standard PDB nomenclature from C*α* through the *β, γ, δ, ϵ, ζ*, and *η* positions (Figure 1a). An absent atom occupies the coordinates of its immediate parent rather than collapsing onto C*α*. The resulting occupancy pattern distinguishes the twenty amino acids after generation, so sequence can be recovered without a separate classification head.

**Figure 1.**
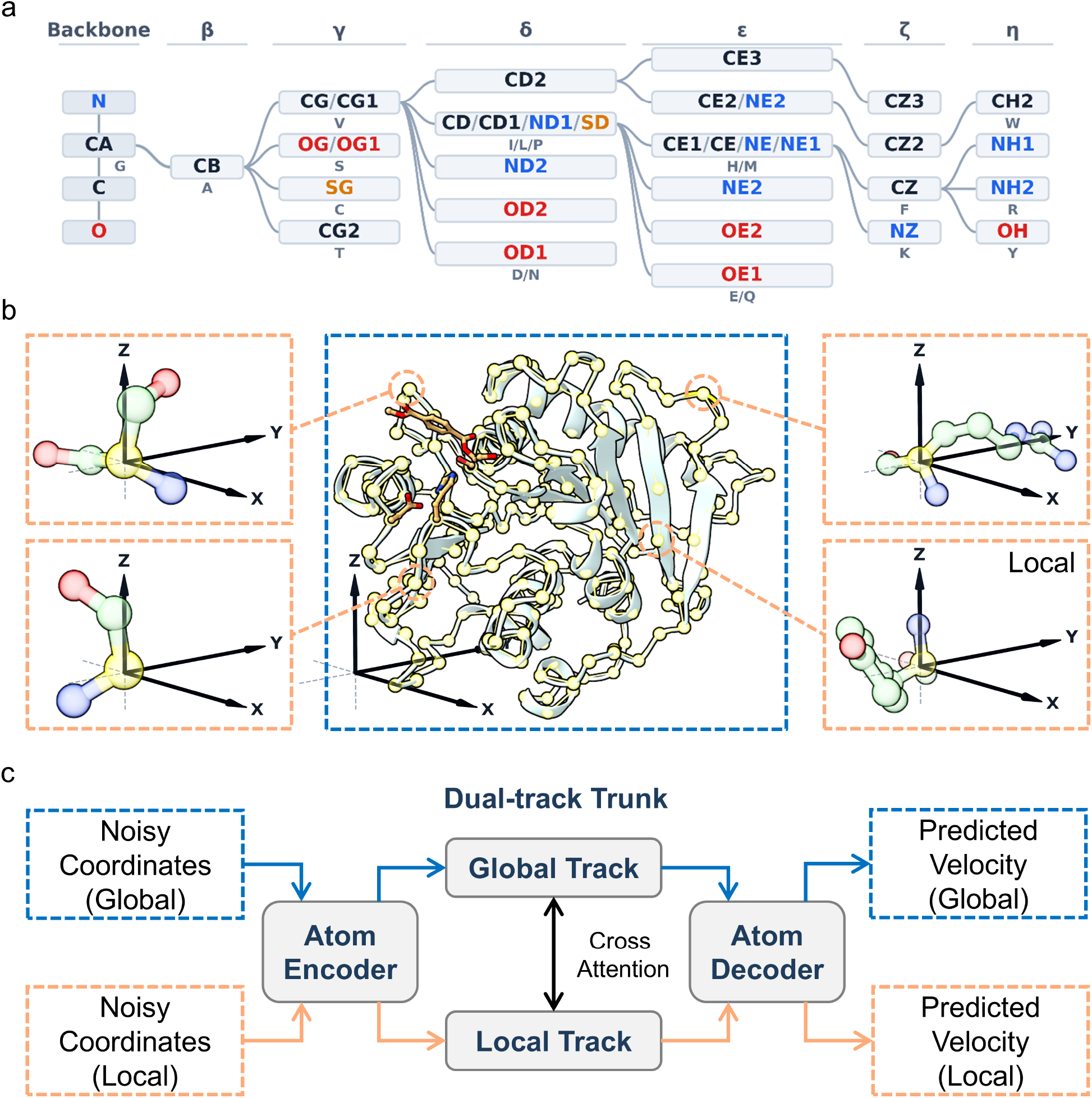
A nested-tree, 28-atom representation and global-local coordinate frame. (a) Atom ordering following standard PDB nomenclature, connected outward from C*α* through the *β, γ, δ, ϵ, ζ*, and *η* positions. Atoms absent from a residue occupy the coordinates of their immediate parent, producing a residue-specific occupancy pattern. (b) Global and local coordinate frames. C*α* is represented in global coordinates, whereas the remaining atoms are represented in local frames centered on their parent residues. (c) Dual-track denoising architecture. The noisy all-atom state is separated into global and local coordinates and encoded into two tracks. The tracks exchange information through bidirectional cross-attention and are then decoded to predict the velocity fields used to update the noisy coordinates.

Intra-residue atoms are dense, so treating every atom as an independent global token is inefficient. C*α* is therefore denoised in global coordinates, while the remaining atoms are represented in a parent-centered local frame (Figure 1b). For nucleic acids the phosphate atom is the global anchor, and for ligands each atom can serve as its own anchor. A dual-track network encodes the two streams, couples them by bidirectional cross-attention, and decodes velocity fields for the flow-matching sampler (Figure 1c).

### Tangential coefficient-preserving sampling enables robust design success in all-atom protein generation

Flow-matching models can be sampled with either deterministic ODE integration or stochastic SDE integration. Stochastic sampling can improve generation by providing corrections to errors accumulated during numerical integration (Song et al., 2020; Karras et al., 2022; Cao et al., 2023). In PANDA, ODE sampling on both tracks produced structures with poor overall quality (Figure 2a). We therefore evaluated coefficient-preserving sampling (CPS), an SDE formulation introduced in Flow-CPS (Wang and Yu, 2025). CPS improved backbone organization, but structures generated with CPS on both tracks showed severe side-chain collapse, particularly at the protein surface, with many affected side chains adopting alanine-like short configurations (Figure 2b).

**Figure 2.**
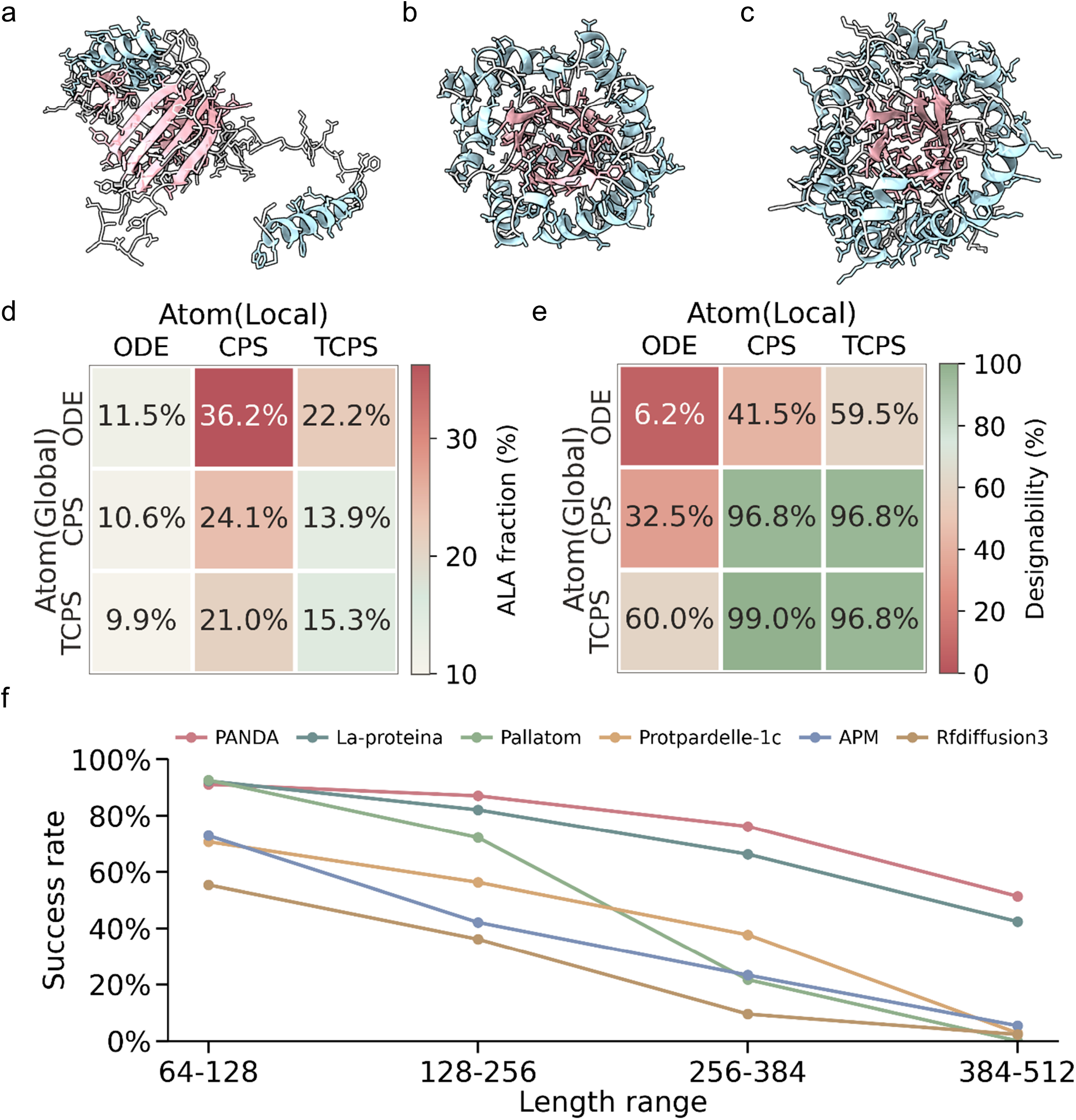
TCPS improves stochastic all-atom sampling. (a) Representative structure generated with ODE on both tracks, showing poor overall structural quality. (b) Representative structure generated with CPS on both tracks, showing an organized backbone but substantial surface side-chain collapse. (c) Representative structure generated with TCPS on both tracks, showing an organized backbone and extended side chains. (d) Alanine fraction across all nine combinations of global and local samplers. The strongest enrichment occurs when CPS is used on the local track, while global sampling modulates the residual alanine fraction. (e) Backbone designability across the same sampler combinations. Designability increases primarily with stochastic sampling on the global track, whereas alanine enrichment is strongest for local CPS. (f) Unconditional design success for proteins from 64 to 512 residues using dual-track TCPS compared with the evaluated all-atom generative models.

The failure is associated with the way isotropic stochastic updates act on the nested-tree local coordinates. Each non-anchor atom is represented relative to its parent, so the distance from the parent contributes to the atomic occupancy and bond-length information used to distinguish residue types. The direction of the corresponding vector determines the orientation of the atom within the local frame. CPS injects stochastic perturbations into all coordinate directions. On the local track, this includes the radial direction associated with the parent-to-child distance. Repeated stochastic replacement along this direction can reduce the distinction between extended and absent side-chain atoms, resulting in enrichment of short side-chain configurations.

We therefore tested a modified stochastic update that retains the correction provided by CPS while excluding the local radial direction. Tangential coefficient-preserving sampling (TCPS) applies the stochastic component in the subspace orthogonal to the local radial vector. The radial component is left to the deterministic denoising update. This modification restored extended side chains while retaining the improved backbone quality obtained with stochastic sampling (Figure 2c).

To separate the contributions of the two tracks, we independently combined ODE, CPS, and TCPS on the global and local tracks. The nine combinations were evaluated for alanine fraction (Figure 2d). The strongest alanine enrichment consistently occurred when CPS was used on the local track, with alanine fractions of 36.2%, 24.1%, and 21.0% for global ODE, CPS, and TCPS, respectively. By comparison, local ODE gave 11.5%, 10.6%, and 9.9%, while local TCPS gave 22.2%, 13.9%, and 15.3%. Thus, the local sampler had the dominant effect on alanine enrichment, although the global sampler also modulated the residual alanine fraction when the local track used TCPS. These results support the view that the collapse is primarily associated with stochastic updates applied along the local coordinate representation rather than with stochastic sampling itself.

Backbone designability showed a distinct dependence on the sampling configuration (Figure 2e). Global ODE produced low designability across local samplers, with success rates of 6.2%, 41.5%, and 59.5% for local ODE, CPS, and TCPS, respectively. Introducing stochastic sampling on the global track increased designability substantially. Global CPS and TCPS reached 32.5% and 60.0% with local ODE, while combinations with local CPS or TCPS reached 96.8% or higher in several settings. Local CPS therefore improved designability when paired with a stochastic global sampler but simultaneously produced the strongest alanine enrichment. Dual-track TCPS provided a favorable balance, reaching 96.8% designability while limiting alanine enrichment to 15.3%.

We next evaluated unconditional design success for proteins spanning 64 to 512 residues. Dual-track TCPS achieved the highest success rate among the PANDA sampling configurations in every tested length range and exceeded the evaluated all-atom generative models across all four ranges (Figure 2f).

### Motif- and ligand-conditioned generation yields atomically precise scaffolds

Functional protein design often requires a generated scaffold to preserve the relative arrangement of catalytic or binding atoms. Existing all-atom scaffolding methods commonly keep motif or ligand coordinates fixed during denoising (Krishna et al., 2024; Wang et al., 2026; Butcher et al., 2025). PANDA instead conditions on pairwise distances among motif atoms, so those atoms remain free to move with the scaffold while their internal geometry is retained (Figure 3a). Pairwise distances do not uniquely determine chirality, so an additional sampling correction is used to resolve the mirror-image ambiguity. For ligand conditioning, PANDA can optionally use relative accessible surface area (RASA) to control the burial of individual ligand atoms.

**Figure 3.**
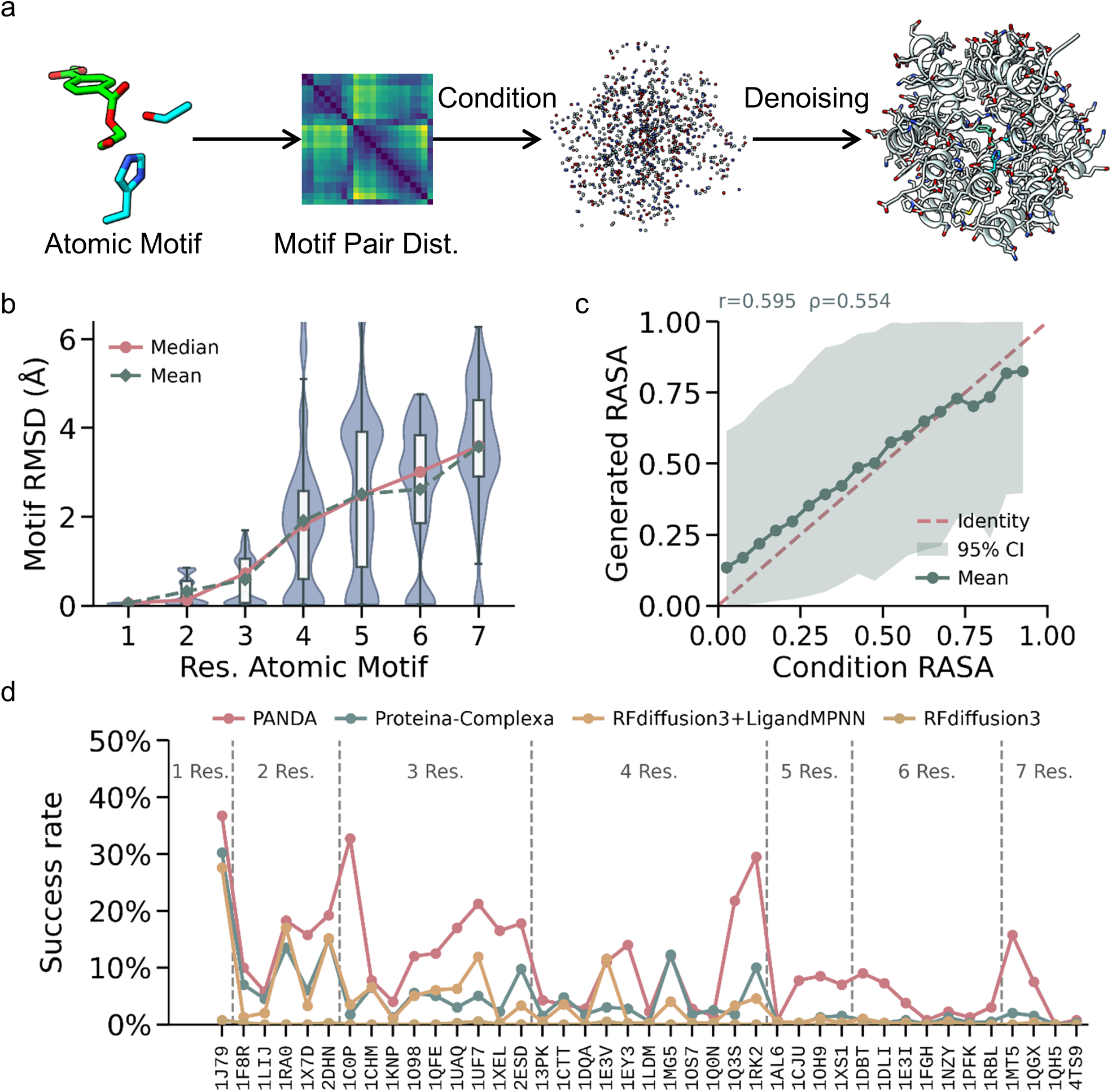
Atomic motifs and ligands condition all-atom generation without coordinate freezing. (a) Functional atomic motifs are converted into pairwise distance matrices that condition denoising of the complete all-atom structure. (b) Motif RMSD after alignment to preset coordinates, grouped by motif size. RMSD increases and the distributions broaden with motif size. (c) Generated versus prescribed RASA under optional ligand-burial conditioning. Generated burial follows the prescribed direction but occupies a narrower range and regresses toward intermediate values at the extremes. (d) Success rate on the AME benchmark by motif size. PANDA is compared with Proteina-Complexa, RFdiffusion3 with LigandMPNN, and RFdiffusion3.

We first examined motif-conditioned generation independently of complete scaffold quality. Functional motifs were supplied as disconnected atomic fragments and converted to a pairwise distance matrix used during denoising. Motif fidelity was quantified by aligning the generated motif atoms to their preset coordinates and measuring heavy-atom RMSD (Figure 3b). Median RMSD was close to zero for one- and two-residue motifs and increased progressively with motif size, reaching approximately 3.5 Å for seven-residue motifs. The distributions also broadened as motif size increased, indicating greater variability among generated arrangements. This trend is expected because larger motifs retain more degrees of freedom that are not uniquely determined by the pairwise constraints. The results nevertheless show that pairwise-distance conditioning preserves the motif geometry at an angstrom-scale level across the tested motif sizes.

We next evaluated ligand burial through RASA conditioning (Figure 3c). Generated RASA showed a positive association with the prescribed values across the tested range, with Pearson correlation *r* = 0.595 and Spearman correlation *ρ* = 0.554. The generated values, however, occupied a narrower range than the prescribed values and were biased toward intermediate burial. In particular, highly exposed targets produced generated RASA values above zero, whereas highly buried targets produced values below the prescribed upper end. Thus, RASA conditioning consistently shifted ligand burial in the requested direction but did not reproduce the full target range precisely.

Finally, we evaluated end-to-end scaffolding on the Atomic Motif Enzyme benchmark, which contains functional motifs ranging from one to seven residues (Figure 3d). PANDA achieved higher success rates than the evaluated baseline methods for most targets and retained measurable success as motif size increased. The advantage was particularly evident for several four-to seven-residue motifs, for which the baseline methods often approached zero success. These results show that the pairwise-distance conditioning strategy can support end-to-end motif scaffolding even when the conditioned motif contains multiple residues.

## Discussion

The core contribution of PANDA is the nested-tree representation, which places residue identity inside the same Cartesian occupancy pattern used for structure rather than in a separate discrete head. Dual-track denoising and TCPS are adaptations of this representation. Pairwise-distance conditioning extends it to functional motifs without freezing atom coordinates.

Sampler ablation shows that stochastic updates must respect the local radial geometry of the nested tree. Local CPS produces the strongest alanine enrichment, whereas dual-track TCPS balances designability against this collapse. Sampling schemes therefore cannot be transferred unchanged from backbone-only models.

Pairwise-distance conditioning retains motif geometry at angstrom scale for motifs of up to seven residues, although RMSD increases with motif size. RASA shifts ligand burial in the requested direction but does not cover the full prescribed range.

Several limitations should be considered. Self-consistency design success is evaluated using a structure predictor and therefore measures whether generated sequences and structures are mutually compatible rather than thermodynamic stability, solubility, or biochemical activity. Motif RMSD and AME success assess geometric scaffolding and do not establish catalytic function. Although the representation supports nucleic acids and ligands, the systematic evaluations reported here focus primarily on protein generation. The residual dependence of alanine frequency on global stochasticity also indicates that the global and local tracks are not completely independent because local frames evolve with the global structure. Finally, RASA conditioning does not yet provide precise control at the extremes of ligand burial.

Overall, PANDA shows that residue identity and all-atom structure can be generated as one continuous coordinate process. Further improvements in sampling, functional conditioning, and experimental validation should clarify the extent to which this framework can support the design of proteins with specific biochemical functions.

## Methods

### Model and generative objective

PANDA uses conditional flow matching to predict velocity fields for global and local atomic coordinates. Each residue is represented by 28 nested-tree slots. C*α* is denoised in global coordinates, while the remaining atoms are represented in a C*α*-centered local frame. The denoiser contains separate global and local atom encoders, a dual-track trunk with bidirectional cross-attention, and separate atom decoders. Noisy coordinates are obtained by interpolation between Gaussian priors and clean coordinates, with separate noise scales and coupled global and local time variables. Further architectural details are provided in Supplementary Information, Section 2.

### Training

Training examples were obtained from PDB-REDO refined experimental structures and the ESMFold structural atlas. Protein chains of 32–384 residues were retained after the filtering procedures described in Supplementary Information, Section 1. Structures were clustered using Foldseek 3Di linclust into a single length-384 catalogue, and training entries were sampled with weights computed from cluster size, entry type and component composition. The flow-matching objective uses mask-normalized velocity MSE together with SmoothLDDT and late-stage coordinate and distance losses. Optimization uses AdamW with BF16 training and an exponential moving average checkpoint.

### Sampling

Inference uses TCPS with 400 integration steps. For each step, the denoiser predicts global and local velocities. The deterministic update follows the predicted velocity, while the stochastic correction is projected onto the subspace orthogonal to the predicted clean local vector. The default stochasticities are *η*_G_ = 0.7 and *η*_L_ = 1.0. ODE, CPS and TCPS are also evaluated independently on the two tracks for the sampler ablation.

### Functional conditioning

Motif conditioning uses the pairwise distance matrix among selected motif atoms. Condition atoms remain unfrozen during denoising. A chirality correction is applied during sampling because pairwise distances permit mirror-image configurations. RASA is an optional ligand-level conditioning feature that is supplied directly to the denoiser when specified. Unconditional sampling does not use RASA. Further details are provided in Supplementary Information, Section 4.

### Evaluation

Unconditional design success is measured by recovering the sequence from generated all-atom coordinates, predicting the corresponding structure with ESMFold (Lin et al., 2023), and requiring C*α* self-consistency RMSD below 2 Å and pLDDT above 75. Backbone designability uses ProteinMPNN (Dauparas et al., 2022) followed by ESMFold. Unless otherwise stated, 1,000 structures were sampled for each length range and 100 samples were generated per AME target per length range. Further evaluation and baseline settings are provided in Supplementary Information, Section 5.

## Acknowledgements

This work was supported by the National Key Research and Development Program of China (No. 2023YFA0914403), the National Natural Science Foundation of China (U23A20219), and the Anhui Provincial Science and Technology Innovation Breakthrough Project (No. 202423110050046).

This work received computational resources support from the Intelligent Computing Platform at the Tianjin Institute of Industrial Biotechnology, Chinese Academy of Sciences.

## Code availability

Code for PANDA will be released at https://github.com/Jay-baijie/PANDA.

## Supplementary information

**Figure S1.**
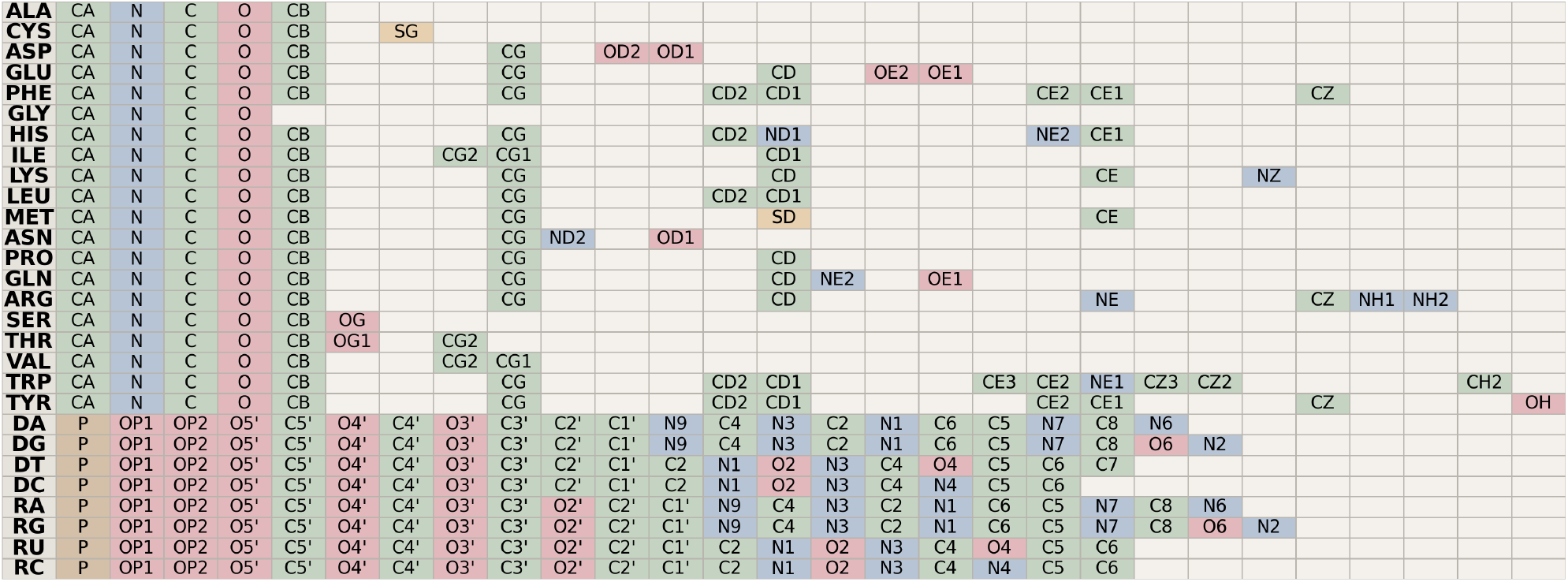
Nested-tree atom-slot representation. Twenty-eight atomic slots are arranged from the global anchor through the nested atom hierarchy. Absent atoms inherit the coordinates of their parent.

### 1 Data preparation and sampling weights

#### 1.1 All-atom representation

Each residue is represented by 28 atomic slots arranged as a nested tree rooted at C*α*. The remaining slots comprise N, C, O and the side-chain atoms ordered by the tree. An atom absent from a residue occupies the coordinate of its immediate parent. The resulting occupancy pattern specifies the twenty canonical amino-acid types and is used to recover sequence after generation. C*α* forms the global track, while the remaining 27 atoms are represented in the C*α*-centred local frame. Nucleic-acid and ligand components use the corresponding anchor definitions.

#### 1.2 Training corpora and filtering

Experimental structures were obtained from PDB-REDO refined coordinates. Entries without reported resolution or with resolution above 3.0 Å were discarded. Protein chains outside 32–384 residues, chains containing unknown residues, and chains with consecutive C*α* distances above 10 Å were removed. Near-terminal polyhistidine tags were trimmed. Chains with incomplete residue masks were retained only when an adjacent ligand interface was present and the remaining backbone satisfied the continuity criterion. At most twenty protein chains were retained per assembly by random subsampling, together with ligands adjacent to the retained polymers. Experimental B-factors were not used as confidence scores.

ESMFold atlas structures were filtered using the confidence criteria provided with the atlas, corresponding to predicted LDDT and predicted TM scores above 0.7. Terminal disordered regions were removed by trimming consecutive terminal residues with fewer than 80 residues remaining in the retained segment, followed by the same length and continuity filters.

#### 1.3 Clustering

Global-atom geometries from the two sources were encoded as 3Di structural sequences and clustered jointly with Foldseek linclust at 50% 3Di identity and 0.8 coverage. The training catalogue used a residue-length upper bound of 384, matching the atom crop budget.

#### 1.4 Training entries and weight-file construction

After clustering, training weights were computed for the length-384 catalogue. For every cluster *C* with size *n*_*c*_, all member sequences were traversed and their corresponding structure files were read. Each valid protein chain generated a single-chain training entry. In addition, each protein chain generated a protein neighbour interaction entry for every annotated neighbouring chain, including protein–protein, protein–nucleic-acid and protein–ligand interactions.

For entry *i*, let *n*_p,*i*_, *n*_n,*i*_ and *n*_l,*i*_ denote the numbers of protein, nucleic-acid and ligand chains in the entry. We define the component-type multiplier as

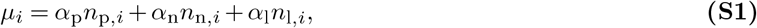

where *α*_p_ = 3, *α*_n_ = 4 and *α*_l_ = 5. The raw sampling weight is

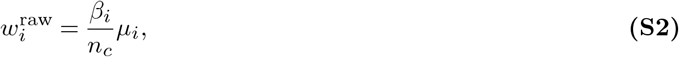

where *β*_*i*_ = 1.0 for a single-chain entry and *β*_*i*_ = 2.0 for an interaction entry. Division by *n*_*c*_ reduces the contribution of structurally redundant clusters, so that individual entries from larger clusters receive lower relative weight.

The entries and their raw weights were collected into the length-384 weight file. Weights were L1-normalised over all valid entries,

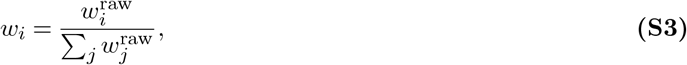

and the resulting categorical distribution was used for sampling with replacement during training. The construction procedure is summarised in Algorithm S1.

### 2 Model architecture

#### 2.1 Network overview

PANDA is a dual-track flow-matching denoiser with approximately 630 million parameters. Static atom and pair features are initialized once per example. Noisy global and local coordinates are encoded separately, processed by a shared dual-track trunk, and decoded into global and local velocity fields. Residue single and pair channels have widths *c*_*s*_ = 768 and *c*_*z*_ = 128, and atom and atom-pair channels have widths *c*_atom_ = 128 and *c*_atompair_ = 16. The trunk contains 16 dual-track blocks. Residue-level attention uses 12 heads and atom-level attention uses 4 heads. Forward computation uses BF16. The complete forward pass is given in Algorithm S4.

#### 2.2 Feature initialization

Atom features include atom indices, element types, charges, reference coordinates, bond information, aromaticity and template masks. Pair features include relative sequence position, reference distances and bond information. Motif distance features are inserted only for selected condition atoms. RASA is encoded as an optional conditioning feature when supplied.

#### 2.3 Atom Encoder, Dual-track Trunk and Atom Decoder

The atom encoder embeds the current noisy coordinates and applies three rounds of pair-biased attention to global atoms and three rounds of axial attention within each residue on local atoms. The resulting atom features are pooled to residue-level representations. Each dual-track block performs bidirectional cross-attention between the global and local representations, updates pair features from both tracks, and applies pair-biased self-attention on each track. The decoder broadcasts the residue representations back to atoms, applies the corresponding atom-level attention blocks, and projects to Cartesian velocities. The main operators are detailed in Algorithms S7–S9 and Algorithms S10–S16.

### 3 Training

#### 3.1 Flow-matching construction

For each training example, the target all-atom coordinates are separated into the global C*α* coordinates 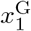 and local coordinates 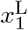. Independent Gaussian priors are sampled as

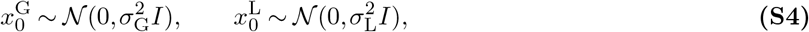

with *σ*_G_ = 10 and *σ*_L_ = 1.5. The global time is sampled by

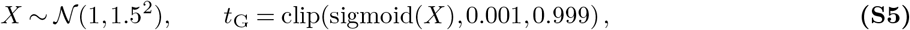

followed by

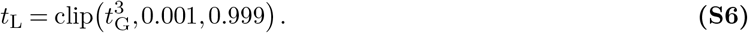

For *k* ∈ *{*G, L*}*, the linear flow-matching interpolant and target velocity are

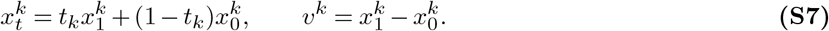

The denoiser predicts 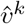 and the corresponding clean-coordinate estimate is

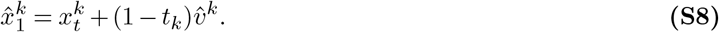

A shared random rotation is applied to the global and local coordinates, with optional translation in the global frame. The same augmentation is applied to the corresponding target and interpolated coordinates. Local coordinates remain expressed relative to their residue anchors.

#### 3.2 Primary velocity losses

Let *m*^G^ and *m*^L^ denote the atom masks after intersecting the atom-validity mask with the noised-atom mask. For each example *b*, the mask-normalised velocity losses are

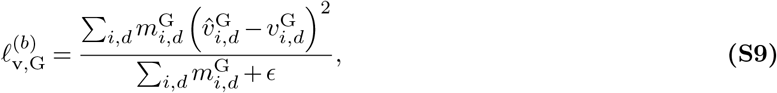

and

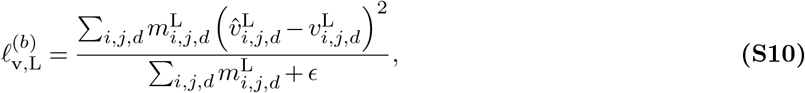

where *ϵ* = 10^−4^. Batch losses are obtained by averaging over examples,

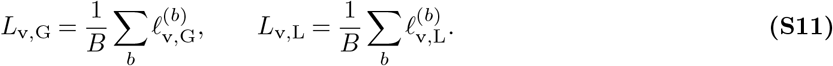

The two terms are divided by 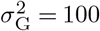 and 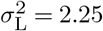, respectively.

#### 3.3 SmoothLDDT loss

A differentiable LDDT-like loss is computed from pairwise distances between predicted and target global coordinates. For a valid residue pair (*i, j*), let

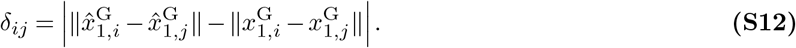

For thresholds *T* = *{*0.5, 1, 2, 4*}* Å, the soft agreement score is

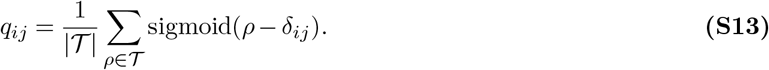

Only target distances below 15 Å are included, together with the upper-triangular pair mask. The resulting loss is

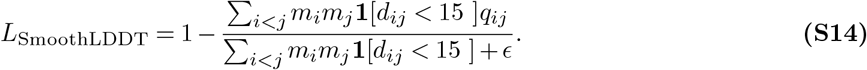

#### 3.4 Late coordinate and pairwise losses

Additional geometric losses are activated only after optimiser step 150,000. The time-weighted coordinate losses are

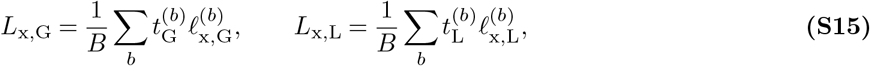

where *ℓ*_x,G_ and *ℓ*_x,L_ use the same mask-normalised form as the velocity losses, replacing 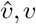 with 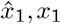. For template-masked global atoms, the template pairwise-distance loss is

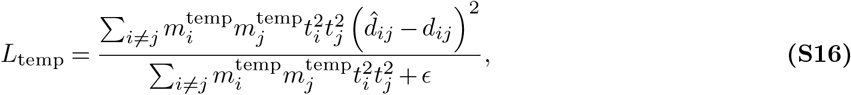

where 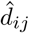 and *d*_*ij*_ are predicted and target global C*α* distances.

For each valid residue, up to eight nearest neighbours are selected according to the target C*α* coordinates. Local coordinates are first transformed to the global frame,

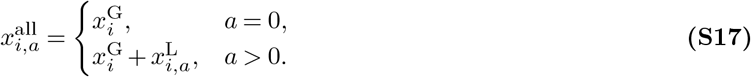

All valid atom pairs between a centre residue and its selected neighbours are then compared. The neighbour loss is

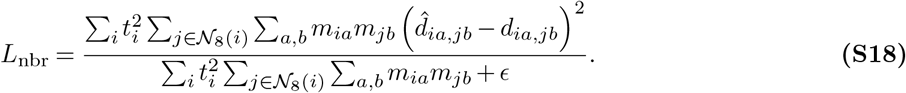

Neighbour identities are determined from the target structure and are not recomputed from predictions. The complete training objective is

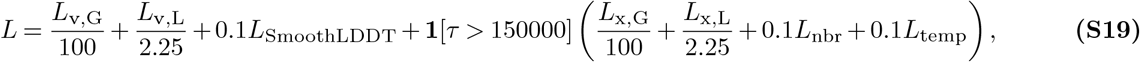

where *τ* denotes the optimiser step. The factors 100 and 2.25 are the squared prior scales.

#### 3.5 Optimisation and parameter averaging

Training uses DeepSpeed with BF16 computation and AdamW. The learning rate is 10^−4^ with *β*_1_ = 0.9, *β*_2_ = 0.999, *ϵ* = 10^−6^ and weight decay 10^−3^. A linear warmup is used for the first 1,000 optimiser steps. The gradient norm is clipped to 10, and the default micro-batch size is 8 structures per GPU. Gradient accumulation is handled by the distributed training engine, with the optimiser step counter advanced at accumulation boundaries.

An exponential moving average is optionally maintained for evaluation. If the EMA decay is *ρ* and the state is updated every *N* optimiser steps, the update is

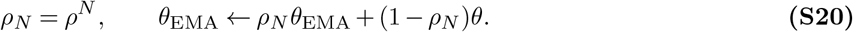

The EMA state is maintained in FP32 on rank 0 and saved separately with the model checkpoint.

### 4 Sampling and conditioning

#### 4.1 Sampling schedule

Inference uses 400 steps by default. The global time grid is constructed from positive normalised gaps and then monotonically warped. The local grid is

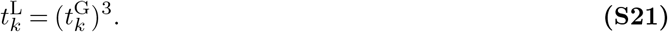

At the final step, the deterministic clean estimate is returned. Local coordinates are converted to the global frame after sampling.

#### 4.2 Tangential coefficient-preserving sampling

Coefficient-preserving sampling modifies the stochastic update by interpolating between the current state and the denoiser-predicted clean state while preserving the prescribed coefficients of the flow-matching trajectory. TCPS further removes the component of the stochastic perturbation parallel to the predicted clean direction before applying the update. This construction retains stochastic correction while restricting the injected perturbation to the tangential subspace defined by the current prediction. The complete update, including the coefficient definitions and tangential projection, is given in Algorithm S17. The default stochasticities are *η*_G_ = 0.7 and *η*_L_ = 1.0.

#### 4.3 Motif distance conditioning

For selected motif atoms, PANDA computes

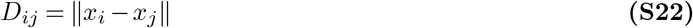

and supplies the pairwise distance matrix as a template pair feature. Condition atoms remain unfrozen during denoising. A late template distance loss penalises deviations between predicted and prescribed motif distances. Because pairwise distances are invariant to reflection, a chirality correction is applied during sampling. Predicted clean condition-atom coordinates are transformed to the global frame and aligned to the template by weighted Kabsch SVD. If the optimal orthogonal transformation is improper, the correction

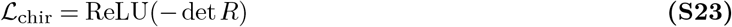

is applied to the predicted velocity with default scale *λ* = 1. The correction is skipped when fewer than four condition atoms are present. The procedure is given in Algorithm S18.

#### 4.4 Optional RASA conditioning

RASA is defined per ligand atom and normalised to [0, 1], with 0 corresponding to buried and 1 to exposed. When a RASA condition is specified, it is encoded directly as an additional conditioning feature. When no RASA condition is supplied, the corresponding feature is absent. RASA is optional and is not used for unconditional sampling.

### 5 Evaluation

#### 5.1 Unconditional generation

Structures were sampled for length ranges 64–128, 128–256, 256–384 and 384–512 residues. Unless otherwise stated, 1,000 structures were generated per range using the EMA checkpoint and 400 TCPS steps.

#### 5.2 Sampler ablation

ODE, CPS and TCPS were crossed independently on the global and local tracks, producing the nine combinations

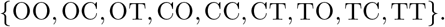

Each combination used 1,000 structures per length range. Alanine fraction and backbone designability were evaluated with the same downstream procedures.

#### 5.3 Design success and designability

For self-consistency, the amino-acid sequence was recovered from generated all-atom coordinates and folded with ESMFold (Lin et al., 2023). C*α* RMSD after Kabsch alignment and pLDDT were calculated. A sample was successful when scRMSD was below 2 Å and pLDDT exceeded 75. Backbone designability used Protein-MPNN (Dauparas et al., 2022) with eight redesign sequences per backbone followed by ESMFold. A backbone was considered designable when at least one redesign satisfied the same thresholds.

#### 5.4 AME benchmark

The Atomic Motif Enzyme benchmark contains functional motifs spanning one to seven residues. Each target was sampled 100 times per length range. Motif fidelity was measured by heavy-atom RMSD after alignment to the preset motif. End-to-end success required motif fidelity together with scaffold self-consistency.

#### 5.5 Baseline methods

Unconditional and AME comparisons used the public checkpoints of Pallatom (Qu et al., 2024), APM (Chen et al., 2025), La-Proteina (Geffner et al., 2025a), Proteina-Complexa (Didi et al., 2026), RFdiffusion3 (Butcher et al., 2025) and ProtPardelle where applicable. Baseline sampling followed the published or official configurations. The settings used for the reported comparisons are summarised in Table S1.

**Table S1.** Sampling settings used for PANDA and baseline methods.

| Method | Checkpoint or configuration | Steps or schedule | Notes |
| --- | --- | --- | --- |
| PANDA | EMA, length-384 run | $T = 400$ TCPS | $\eta_G = 0.7$, $\eta_L = 1.0$ |
| Pallatom | official checkpoint | $T = 200$ | authors' default settings |
| APM | official refine checkpoint | authors' default | unconditional configuration |
| La-Proteina | official unconditional configuration | authors' default | all-atom model |
| Proteina-Complexa | paired AE and diffusion checkpoint | authors' default | matched evaluation protocol |
| RFdiffusion3 | Foundry remapped checkpoint | authors' default | optional LigandMPNN for baseline runs |
| ProtPardelle | cc91 epoch 383 | authors' default | unconditional all-atom configuration |

### 6 Algorithms

The algorithms are organised according to the execution order of data preparation, training, model inference and sampling. They specify the implementation details needed to reproduce the reported procedures.

#### 6.1 Training-weight construction

The following procedure constructs the length-384 weight file used for weighted training-sample selection.

##### Algorithm S1

Construction of the training-weight file

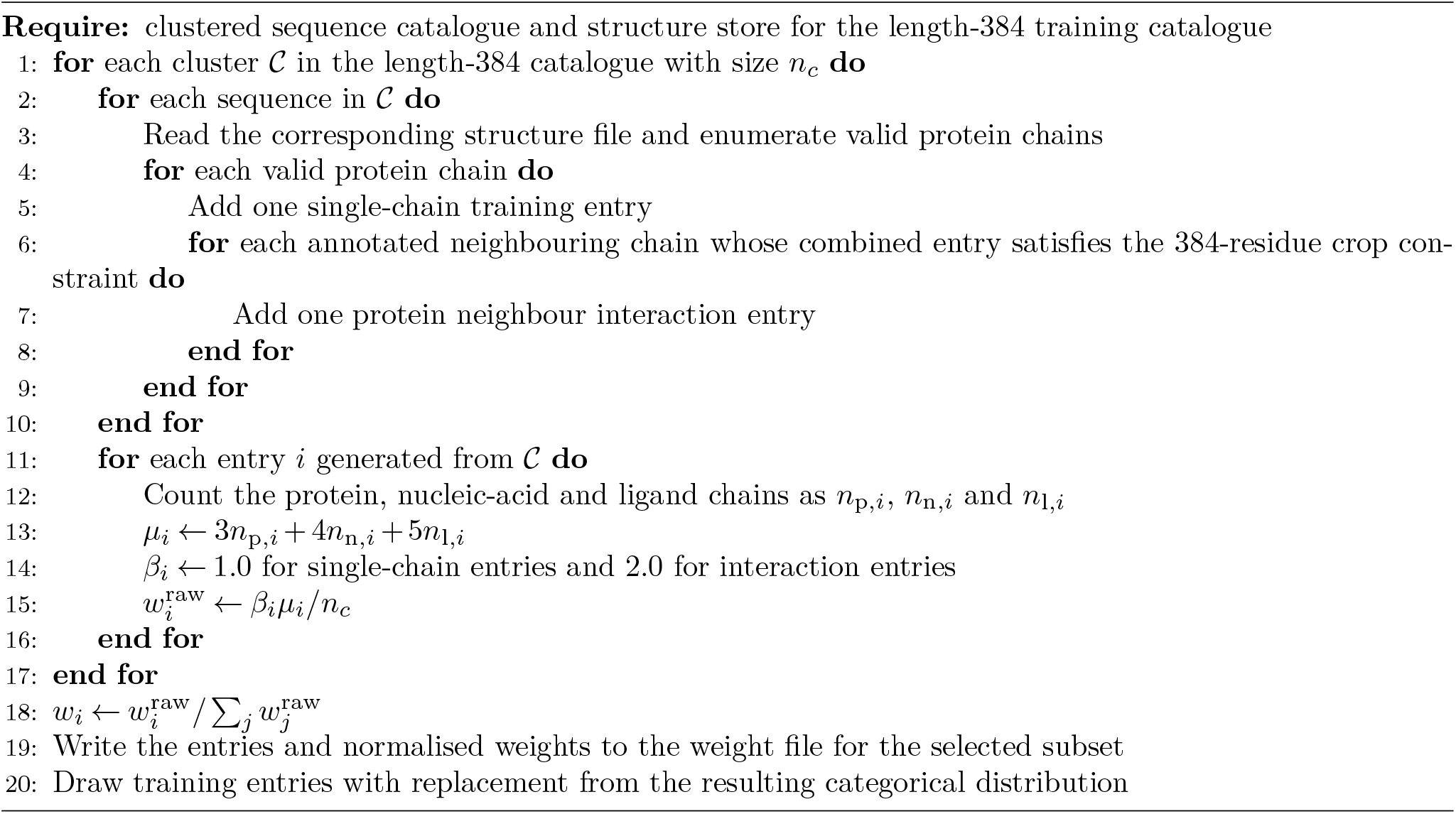

#### 6.2 Training and sampling

##### Algorithm S2

Training step

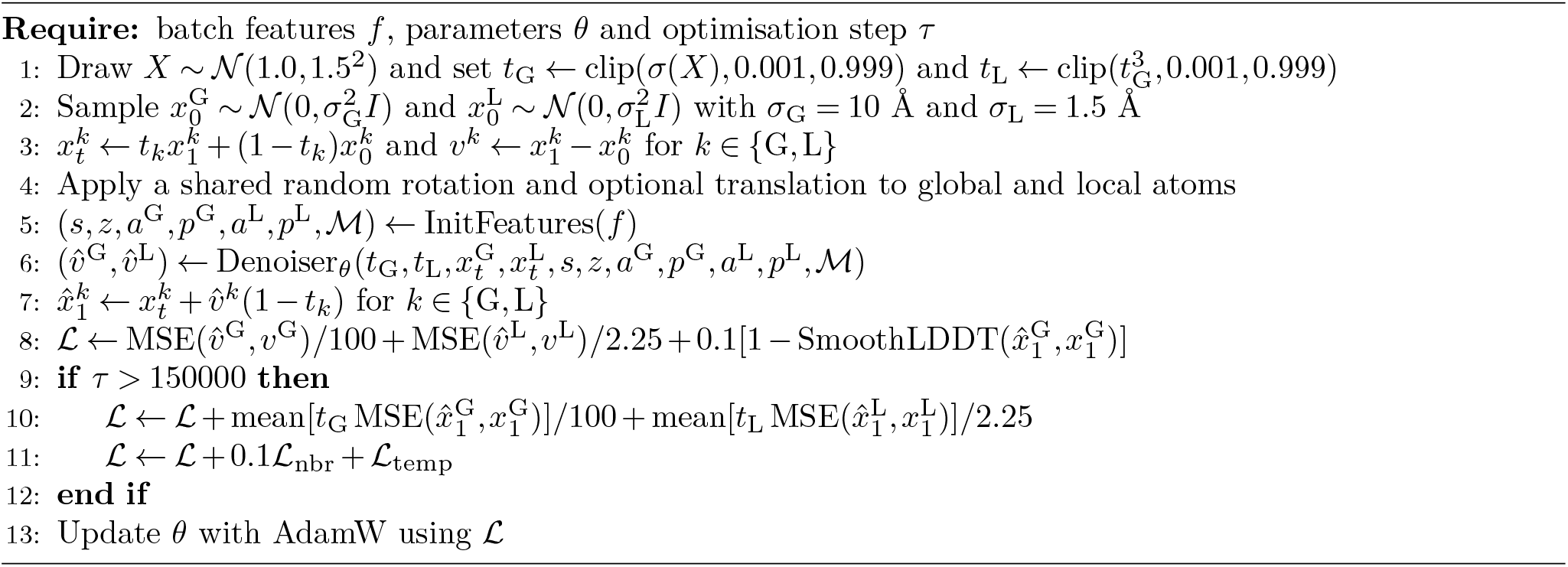

Velocity MSE is mask-normalised. The factors 100 and 2.25 correspond to 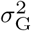 and 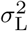. SmoothLDDT uses thresholds {0.5, 1, 2, 4} Å within a 15 Å cutoff. ℒ_nbr_ is an eight-neighbour pairwise-distance loss and ℒ_temp_ is a template pairwise-distance loss.

##### Algorithm S3

TCPS sampling

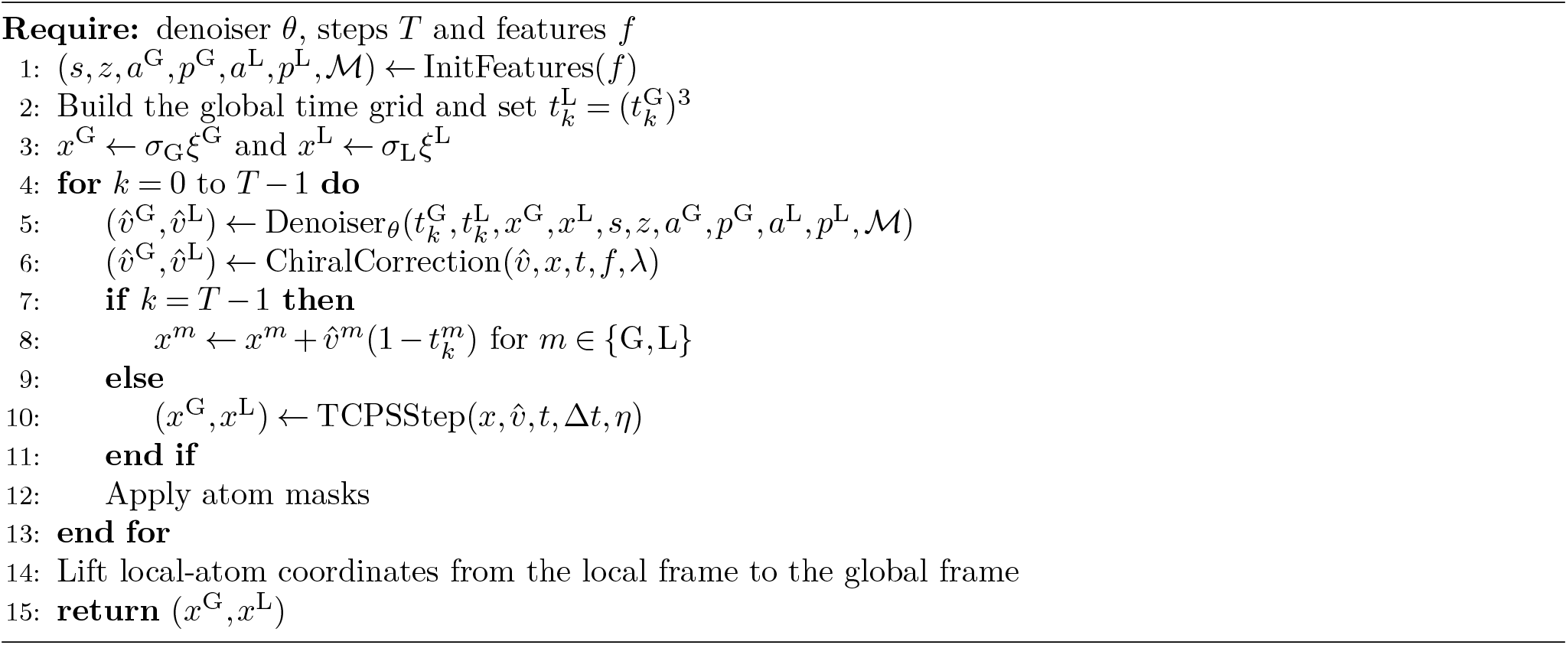

#### 6.3 Denoiser forward pass

##### Algorithm S4

Denoiser forward

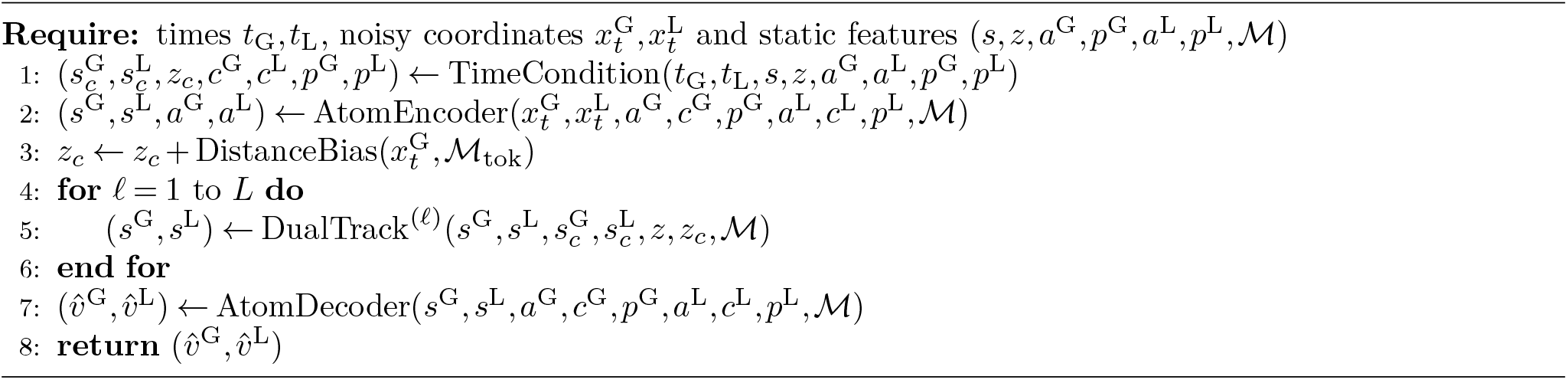

##### Algorithm S5

InitFeatures

Here Φ denotes a Fourier-feature MLP. RelPos embeds signed intra-chain residue offsets with sine and cosine features and assigns zero to cross-chain pairs. Template distance features are gated by the template mask. The RASA embedding is included only when a RASA condition is supplied.

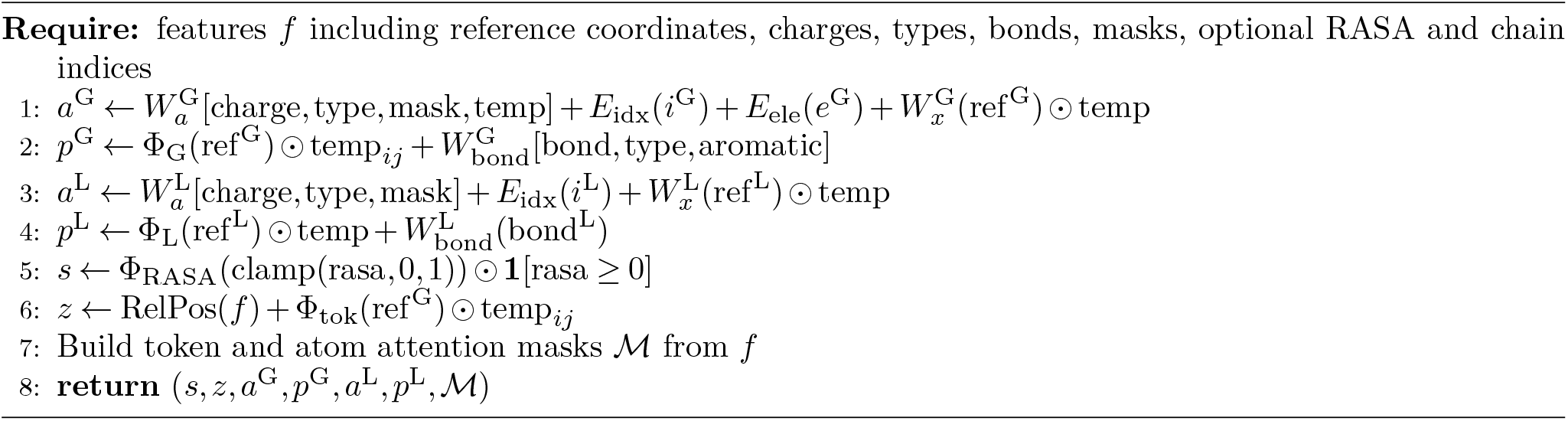

##### Algorithm S6

TimeCondition

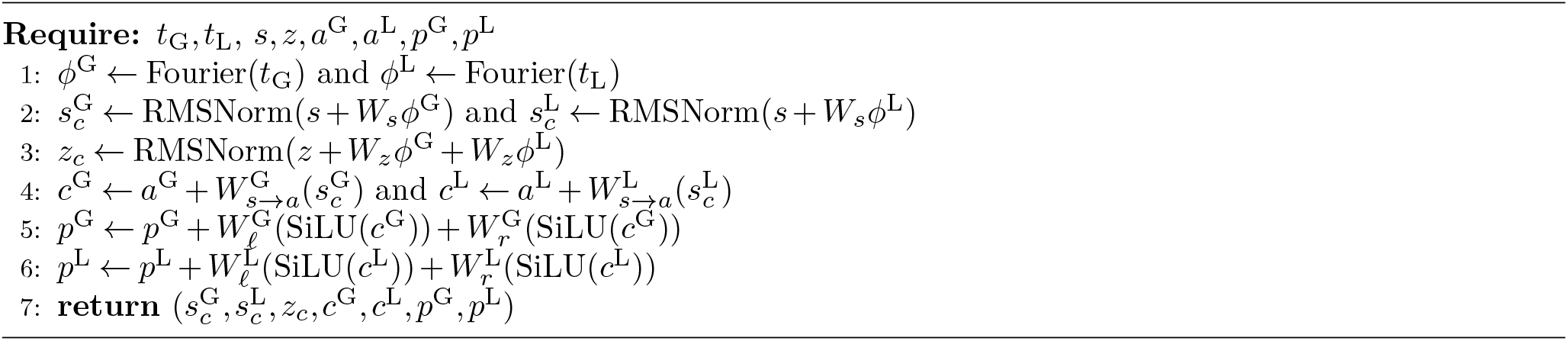

##### Algorithm S7

AtomEncoder

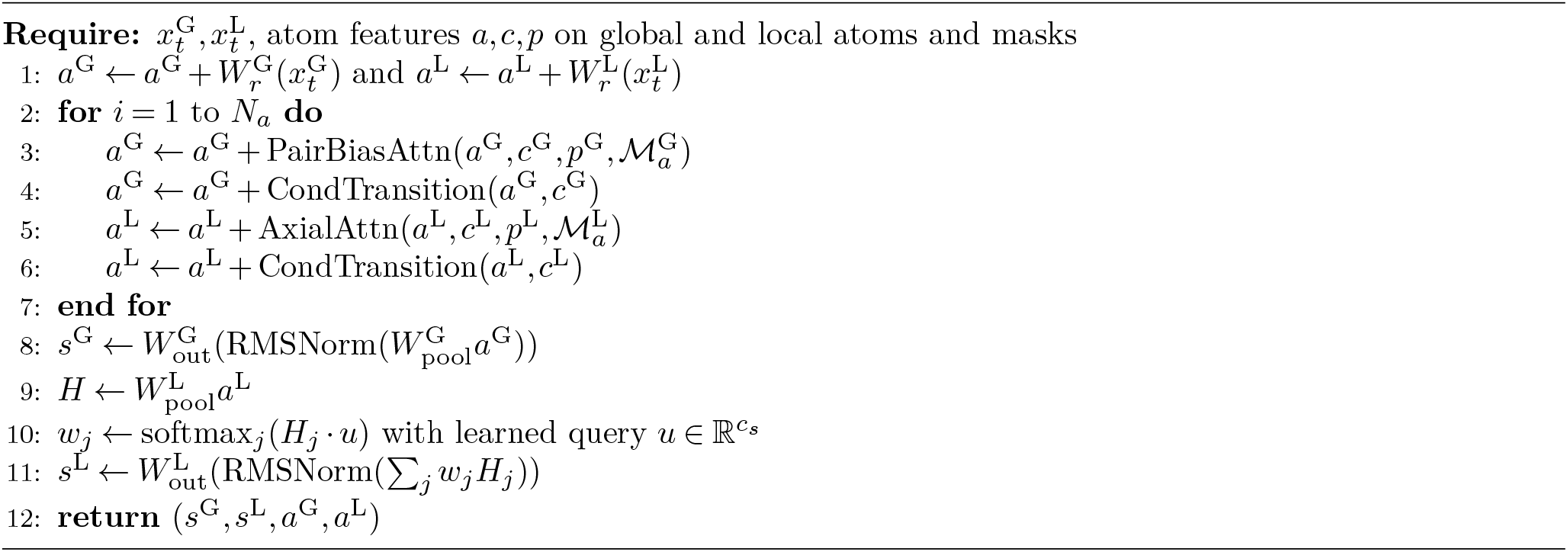

##### Algorithm S8

DualTrack block

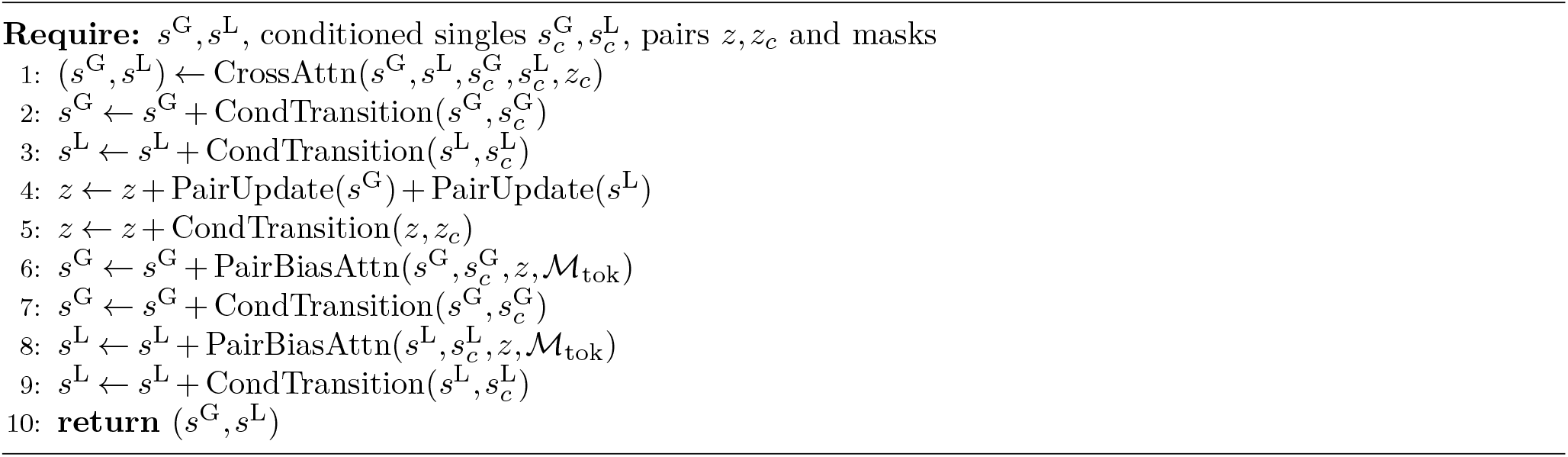

##### Algorithm S9

AtomDecoder

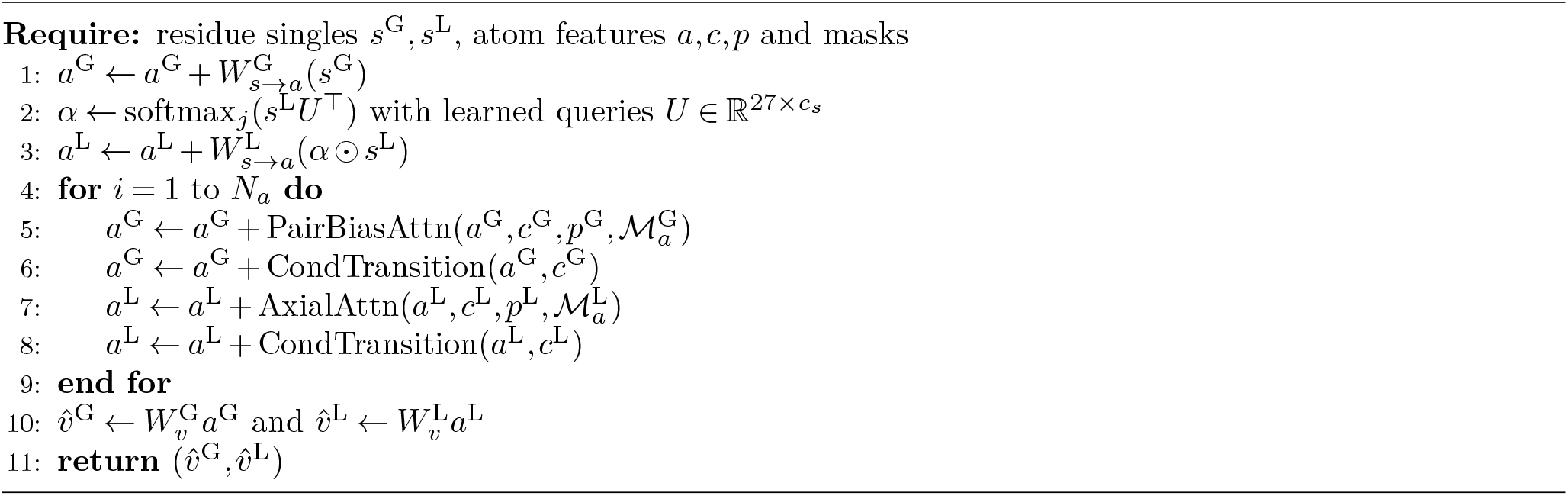

#### 6.4 Shared components

##### Algorithm S10

AdaRN

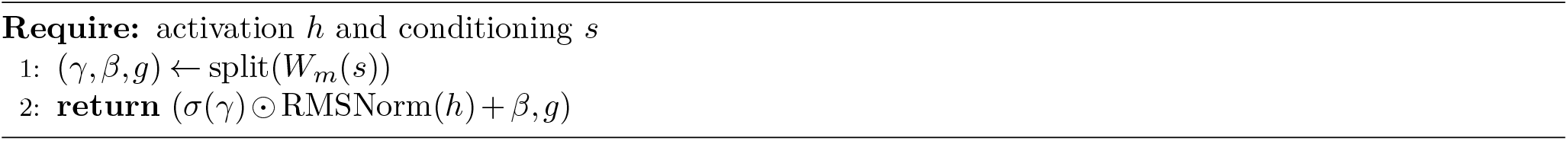

##### Algorithm S11

CondTransition

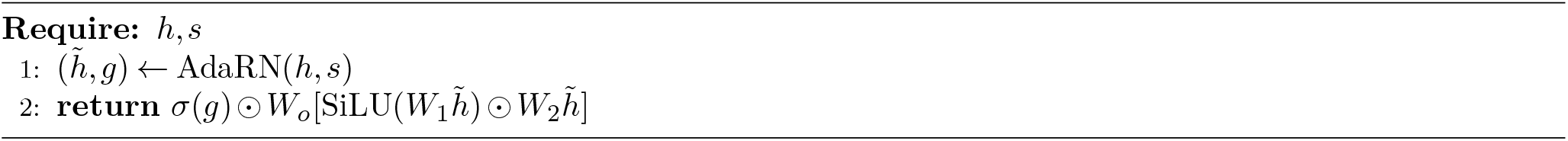

##### Algorithm S12

PairUpdate

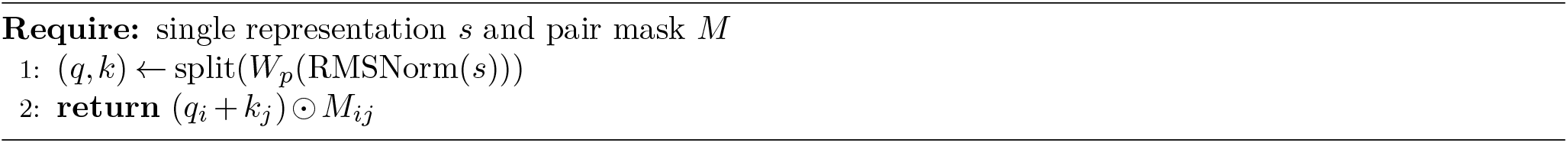

##### Algorithm S13

DistanceBias

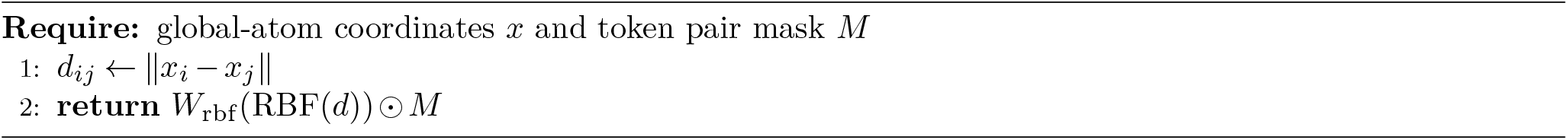

##### Algorithm S14

PairBiasAttn

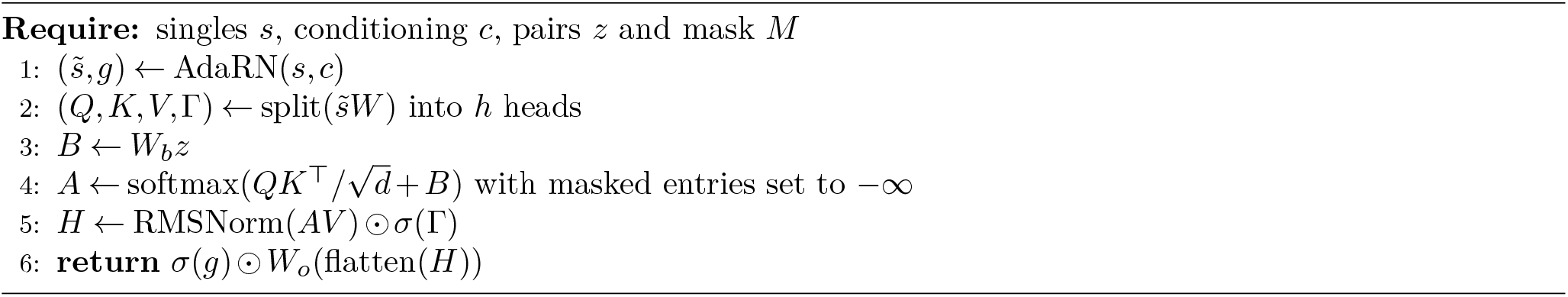

##### Algorithm S15

AxialAttn

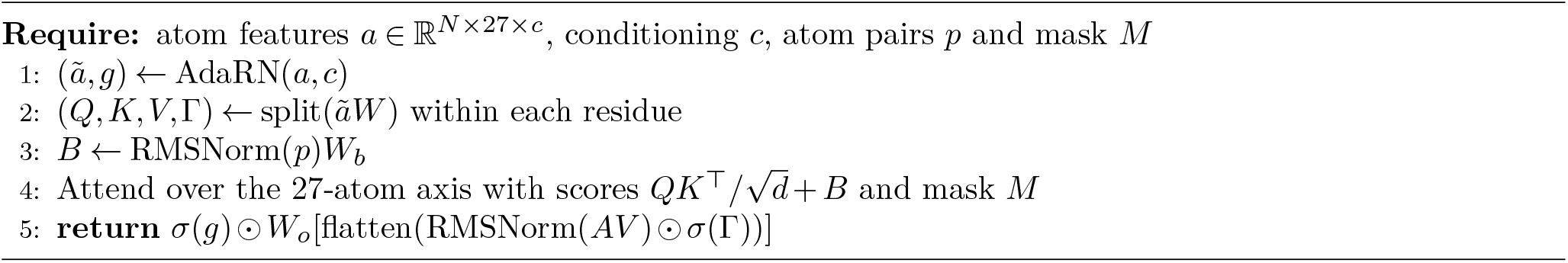

##### Algorithm S16

CrossAttn

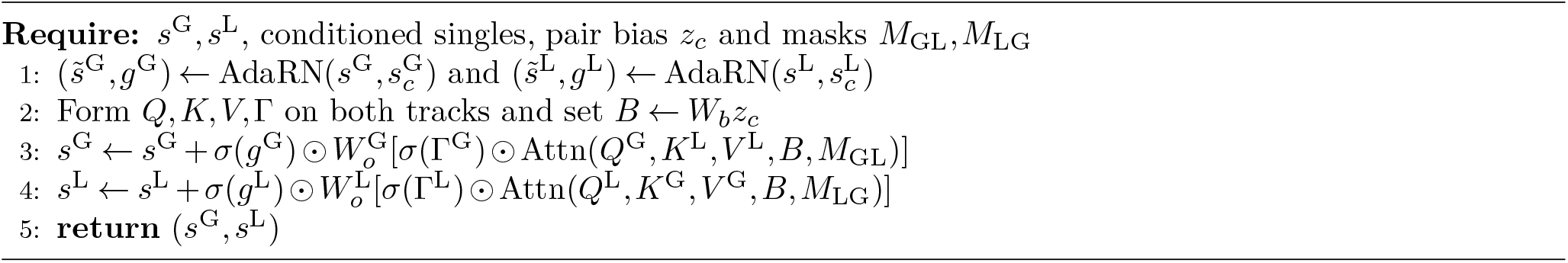

#### 6.5 TCPS update

Let *t*^*′*^ = *t* +Δ*t, η* ∈ [0, 1], *c* = cos(*ηπ/*2), *s* = sin(*ηπ/*2), 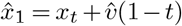 and 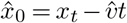. Coefficient-preserving replacement gives

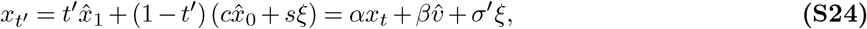

with *α* = *t*^*′*^ + (1 − *t*^*′*^)*c, β* = Δ*t* + (1 − *t*^*′*^)*t*(1 − *c*) and *σ*^*′*^ = (1 − *t*^*′*^)*s*. Tangential projection removes the component along the predicted clean direction,

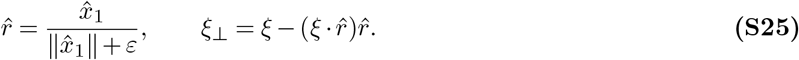

The stochastic term is replaced by *ξ*_⊥_ in TCPS. Default stochasticities are *η*_G_ = 0.7 and *η*_L_ = 1.0.

##### Algorithm S17

TCPS Step

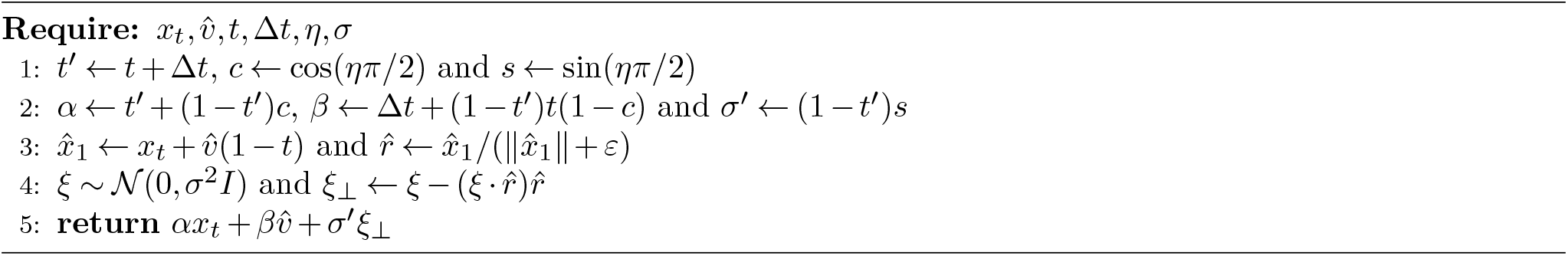

##### Algorithm S18

Chiral correction

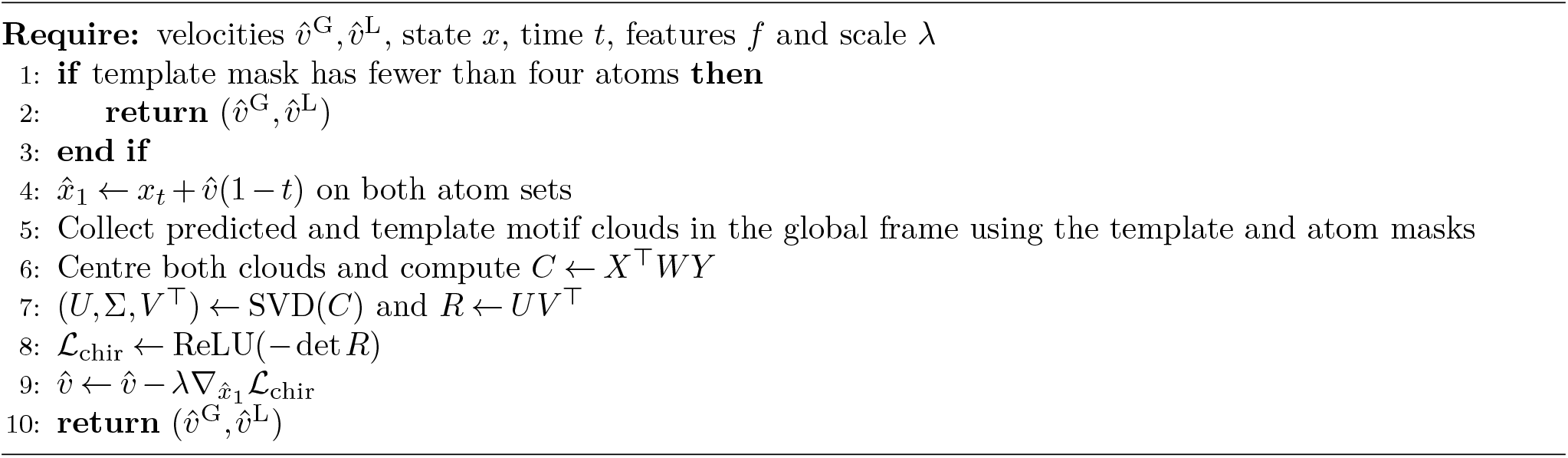

